# Including biotic factors does not improve species distribution models across three trophic levels

**DOI:** 10.1101/2025.08.07.668510

**Authors:** Emelia O Kusi, Catherine Hulshof, Karen Kester

**Affiliations:** Virginia Commonwealth University

**Keywords:** species distribution modeling, *Manduca sexta*, *Manduca quinquemaculata*, *Cotesia congregata*

## Abstract

1. Species distribution is broadly influenced by abiotic factors such as elevation and seasonal variations in temperature and precipitation. Further, for specialist herbivores like *Manduca sexta* and *M. quinquemaculata*, biotic factors, including host plant availability and the presence of a shared parasitoid *Cotesia congregata*, may play a critical role in shaping their spatial patterns. Yet, existing predictive models often focus solely on abiotic factors, with the contribution of biotic factors largely unknown.
2. In this study, we assessed whether abiotic and biotic factors as isolated or combined components of species distribution models (SDM) better predict the distribution of the congeners, *M. sexta* and *M. quinquemaculata*, and their shared endoparasitoid*, C. congregata.* We further evaluated the influence of bioclimate variables and elevation on cultivated and wild host plant distributions of *M. sexta* and *M. quinquemaculata*. To ensure robustness of our SDMs, we constructed them using an ensemble of three predictive algorithms: generalized linear model, maximum entropy, and random forest. We also applied Area Under the Curve (AUC) and True Skill Statistic (TSS) metrics to assess how well each model predicted outcomes. Then, to compare the predictive power between models that used isolated and combined abiotic and biotic factors, we analyzed the values obtained from the AUC and TSS metrics using a one-way ANOVA.
3. Our findings showed that the predictive power of models that used isolated and combined abiotic and biotic factors did not significantly differ. The abiotic-only model identified minimum temperature of the coldest month, elevation, and summer precipitation as the primary determinants of distributions of the *Manduca spp.*, host plants, and *C. congregata,* respectively. Both *Manduca spp.* showed similar responses to minimum temperature of the coldest month and elevation, but response to summer precipitation differed between species. Models accounting for biotic factors only and combination of abiotic and biotic factors revealed cultivated host plants as the most important predictor for the distribution of both the *Manduca spp.* and *C. congregata*. Both *Manduca spp.* And *C. congregata* showed a high likelihood of occurrence with the presence of cultivated host plants. While this finding is not surprising, the responses of *Manduca spp.* to predictors like wild host plants varied between biotic-only and abiotic-biotic models. Similarly, the parasitoid’s response to its host, *Manduca spp.*, also differed between the two models.
4. We therefore suggest that models incorporating either abiotic or biotic factors, or both, can reliably predict species distribution. However, the direction of species responses whether an increase or decrease in likelihood of occurrence may vary depending on the factors included in the model.

## Introduction

The distribution of species underpins the structure of ecosystems, driving biodiversity patterns and informs ecological, conservation, and management strategies (Guisan et al., 2013; Lawler et al., 2010). Every species thrives within a specific range of environmental conditions that supports their occurrence and growth, which defines the limits of their ecological niche (Chase, 2011). These environmental conditions encompass abiotic factors such as climate (Stange & Ayres, 2010; Thomas, 2010) and elevation (Hodkinson, 2005), along with biotic interactions, including top-down effects from predation and parasitism (Giannini et al., 2013; Léandri-Breton & Bêty, 2020), and bottom-up influences such as resource availability and host plant quality (Awmack & Leather, 2002; Gripenberg & Roslin, 2005).

Predicting the geographic range of insects is vital, especially in the context of climate change. As ectotherms, abiotic factors such as temperature, precipitation, and seasonal climatic fluctuations influences insect morphogenesis, reproduction, dispersal, population, and survival (Paaijmans et al., 2013; WallisDeVries et al., 2011). Across North America, various notable insects such as the gypsy moth, *Lymantria dispar* (Régnière et al. 2009) and many species of bark beetles (Bentz et al., 2010; Safranyik et al., 2010) have shown changes in their distribution and impact in response to climate change. Likewise, biotic factors such as host plants and natural enemies affect establishment of ecological niche, population and distribution (Garvey et al., 2020; Giannini et al., 2013). For example, generalist predators like lacewings (Chrysopidae) and ladybird beetles (Coccinellidae) effectively reduce aphid populations, but control is even stronger when specialist natural enemies, such as parasitoid wasps, are present (Diehl et al., 2013; Dixon, 2000).

The distribution and persistence of specialist herbivores and their parasitoids, such as *Manduca sexta* and *M. quinquemaculata* (Lepidoptera: Sphingidae), alongside their shared hymenopterous endoparasitoid, *C. congregata*, are likely to depend on abiotic tolerances, as well as parasitoid and host plant occurrences (Amaya et al., 2005; Contreras et al., 2013; M. A. Garvey et al., 2020; Kingsolver et al., 2021). *Manduca sexta* and *M. quinquemaculata* are lepidopterans whose life cycles illustrate the dual roles of herbivores and pollinators (Bossart & Gage, 1990). During their caterpillar stage, these species exhibit significant herbivory on plants in the Solanaceae family including economically important crops such as tobacco, tomato, and pepper (Yamamoto & Fraenkel, 1960). This stage also exposes them to parasitism by their shared endoparasitoid, *C. congregata;* however, field observations indicate higher preference of this parasitoid for *M. sexta* (Kennedy, 2003; Villanueva, 1998). An adult female *C. congregata* injects eggs into the caterpillar host’s body, where the wasp larvae develop and feed on nutrients in the host hemolymph, eventually leading to death of the host (Villanueva, 1998). When *M. sexta* and *M. quinquemaculata* escape attack by natural enemies and reach adulthood, they transform into pollinators.

Further, wild and cultivated solanaceous plants can support reproductive success and larval development of *Manduca spp*. (Garvey et al., 2020). However, domestication can compromise herbivore defenses to parasitism compared to their wild ancestors (Garvey et al., 2020), which may have repercussion on *Manduca spp*. distribution.

*Manduca sexta* and *M. quinquemaculata* have unique habitats and regions where their distributions overlap (Bossart & Gage, 1990) and exhibit differences in response to thermal conditions (Malinski et al., 2023), yet the mechanisms that regulate their spatial and temporal distribution remain enigmatic. A widely used approach for elucidating the determinants of species distribution is species distribution models (hereafter, SDMs) (Elith & Leathwick, 2009; Guisan & Zimmermann, 2000). SDMs are statistical tools that predict the geographic distribution of species by analyzing the relationship between species occurrence data and environmental variables (Elith & Leathwick, 2009; Guisan & Zimmermann, 2000).

Models built to predict species distributions mostly have been limited to abiotic variables, so the contribution of biotic factors is poorly understood (Guisan & Thuiller, 2005). The use of abiotic factors as the sole predictor in species distribution remains ubiquitous in the current literature due to the difficulties in quantifying the role of biotic factors (Araújo & Guisan, 2006; Godsoe et al., 2017). Foremost, biotic interactions are inherently intricate and can differ dramatically based on environmental conditions and the presence of other species (Kearney & Porter, 2009). Further, the influence of biotic interactions may be overshadowed by environmental gradients, making it difficult to accurately discern how these interactions shape species distributions (Godsoe et al., 2017).

Advancing species distribution models requires refining the species’ ecological niche and enhancing methods for evaluating model performance and predictor importance. To address this need, we utilized an ensemble of SDMs (generalized linear model, maximum entropy, and random forest) to quantify the role of abiotic and biotic factors in explaining the distribution of two congeneric species, *M. sexta* and *M. quinquemaculata*, and their shared parasitoid, *C. congregata*. We further assessed the influence of abiotic factors on the distribution of cultivated and wild host plants of the two *Manduca spp*. By combining robust statistical methods with ecological data, we determined which abiotic and biotic factors best explain the current distribution of each species, and whether the inclusion of biotic factors improved the predictive ability of traditional, abiotic-only SDMs. Here, separate SDMs were constructed using abiotic only, biotic only, and both abiotic and biotic factors combined to better understand the contributions and effects on predictive distribution models.

## Materials and methods

### Study system

Our tritrophic system consisted of the solanaceous specialists, *Manduca sexta* L. (“tobacco hornworm”) and *Manduca quinquemaculata* Haworth (“tomato hornworm”) (Lepidoptera: Sphingidae), their major cultivated and wild host plants, and their shared and sole hymenopterous larval endoparasitoid, *Cotesia congregata* Say (Braconidae) (Figure 1). Distributions of all species in this study were limited to the continental United States. Distributions of the two *Manduca* spp. overlap to some extent; however, *M. sexta* is more abundant in the southern U.S., particularly around the Gulf Coast, while *M. quinquemaculata* is more prevalent in the northern belt of the U.S. (Hodges 1971). The distribution of *Cotesia congregata*, the sole hymenopteran larval endoparasitoid of both hornworm species, spans from the southeastern to the northeastern U.S., with occasional occurrences reported in the western U.S (Illinois Natural History Survey; iNaturalist.org; Molina-Ochoa et al., 2003).

**Figure 1:**
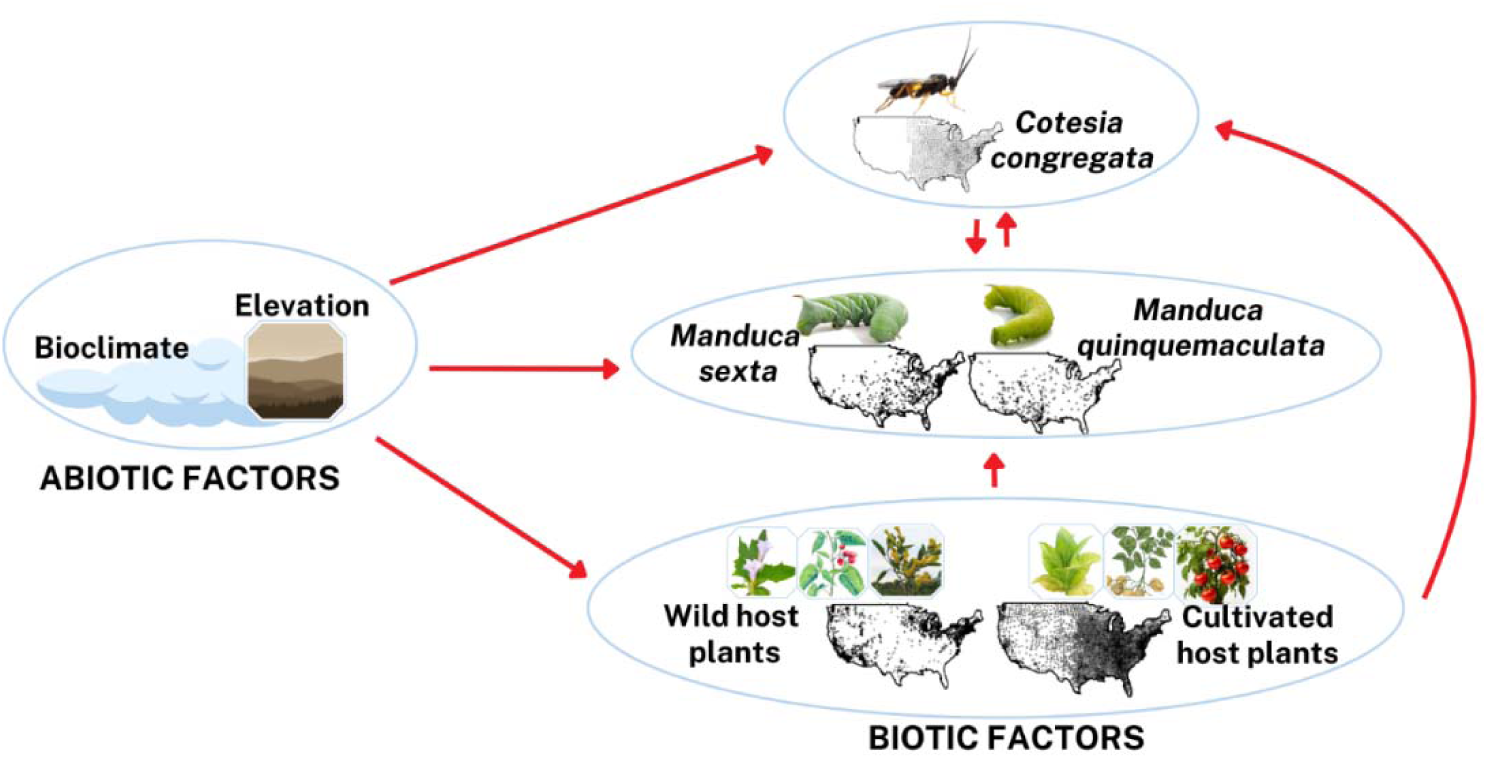
Conceptual diagram of study system. Abiotic factors, specifically bioclimate and elevation, may broadly influence the distribution of all species. Distributions of *M. sexta* and *M. quinquemaculata* may be affected by distributions of their wild and cultivated host plants, as well as distribution of their shared parasitoid, *C. congregata*. Distribution of *C. congregata*, in turn, is likely affected by distributions of hosts, *M. sexta* and *M. quinquemaculata*, and their associated host plants. Red arrows indicate connections between species and predictors.

Plants included both cultivated (non-native) and wild species. Cultivated plants included: *Nicotiana tabacum* L. (cultivated tobacco), *Lycopersicon esculentum* Mill. (*Solanum lycopersicum* L., cultivated tomato), *Solanum tuberosum* L. (potato), *Capsicum annuum* L. (bell peppers, chili peppers), *Solanum dulcamara* L. (bittersweet nightshade) and *Nicotiana alata* Link & Otto (flowering tobacco). Wild species included *Datura wrightii* Regel (sacred datura) and *Solanum eleagnifolium* Cav. (silverleaf nightshade), which are native to the Americas, and *Nicotiana glauca* Graham (tree tobacco); also, being which has naturalized in parts of North America. We examined the distribution of all these species; however, due to insufficient records for some, we focused our analysis on a subset of plant species. Cultivated host plants included *N. tabacum*, *L. esculentum*, and *S. tuberosum*, and wild host plants included *N. glauca*, *D. wrightii*, and *S. dulcamara*. *Capsicum annuum* and *N. alata* were excluded from the analysis despite having sufficient records, as their distributions overlapped significantly with those of other cultivated host plants included in the study. Overall, cultivated host plants exhibited a broader distribution compared to the wild host plants.

### Occurrence records

Occurrence records for the two *Manduca spp.* were compiled from the Centre for Agriculture and Bioscience International (CABI), Integrated Digitized Biocollections (iDigbio), and Global Biodiversity Information Facility (GBIF) using data retrieval package in R (*raster*; Hijmans, 2019). Host plant records were obtained from the United States Department of Agriculture (USDA), CABI, and GBIF. Records for the parasitoid were extracted from GBIF. However, since the available records for the parasitoid were clearly underreported, we supplemented parasitoid occurrences by extracting county-level centroids for all states throughout its range. Duplicates and records older than 1970 were eliminated to match the available climatic data from 1970 onwards. Records were spatially thinned using the R package *spThin* at distances of 1km to reduce the e ects of sampling bias (Aiello-Lammens et al. 2015). Overall, the analytic sample consisted of 1370 records for *M. sexta*, 461 records for *M. quinquemaculata*, and 2845 records for the parasitoid (179 original records and 2666 supplemented records; Figure 2a-c). For wild host plants, a total of 3620 occurrence records were included for *D. wrightii*, *S. dulcamara*, and *N. glauca,* while for the cultivated host plants, a total of 3796 occurrence records were obtained for *N. tabacum*, *S. tuberosum*, and *S. lycopersicum* (Figure 2d-e).

**Figure 2.**
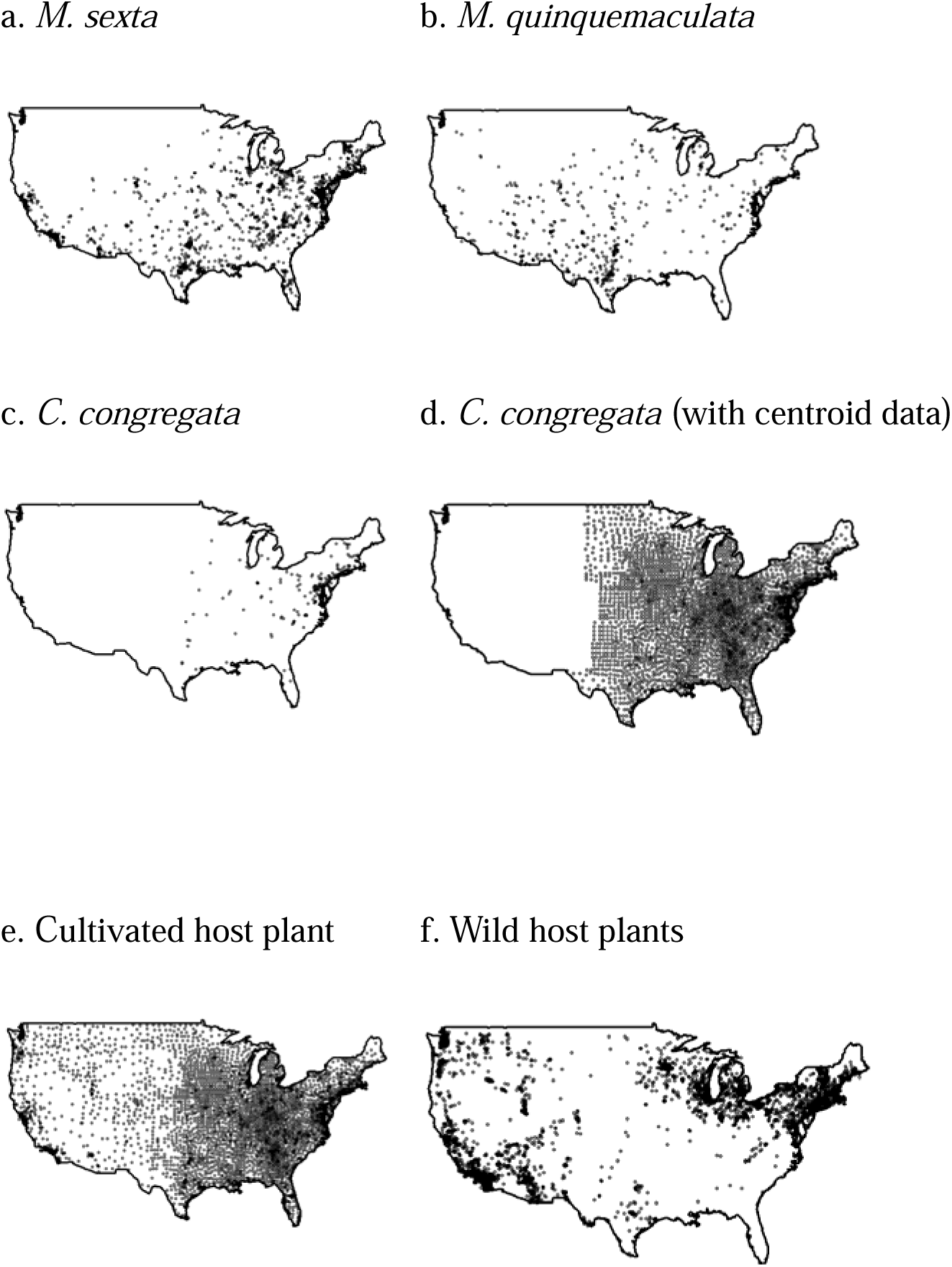
The geographical distribution of species based on occurrence records. Each dot represents a recorded location where the species has been observed except for 2d. which represents the original and supplemental records for the parasitoid, *C. congregata*.

### Abiotic variables

Nineteen bioclimate variables representing annual trends, seasonality and extreme or limiting environmental factors in addition to elevation were downloaded from WorldClim at 2.5 min resolution (Hijmans et al. 2005). These variables included temperature-related metrics such as annual mean temperature, mean diurnal range (the average difference between monthly maximum and minimum temperatures), isothermality (the ratio of mean diurnal range to annual temperature range, scaled by 100), temperature seasonality (standard deviation of monthly temperatures, multiplied by 100), maximum temperature of the warmest month, and minimum temperature of the coldest month. Additional temperature variables included the temperature annual range (difference between the warmest month’s maximum temperature and the coldest month’s minimum temperature) and mean temperatures of the wettest, driest, warmest, and coldest quarters. Precipitation metrics included annual precipitation, precipitation of the wettest and driest months, and precipitation seasonality (coefficient of variation of monthly precipitation values). We also included precipitation totals for the wettest, driest, warmest, and coldest quarters.

A variance inflation factor (VIF) analysis was performed on the abiotic variables using a stepwise procedure from the *usdm* R package (Naimi et al. 2014). To mitigate potential bias and instability in parameter estimation, collinear explanatory variables were excluded from the analysis. The VIF procedure is more precise and the most widely used approach to control for multicollinearity, where variables with a VIF greater than the threshold of 7 were excluded (Chartterjee and Hadi 2006). However, the minimum temperature of the coldest month and precipitation of the driest month were retained due to their biological significance and strong influence on insect survival (Eisen et al. 2015; Pollard 1988; Roy et al. 2008; Vandyk et al. 1996). Thus, a total of nine abiotic variables were included in the SDMs to assess the role of abiotic factors: Maximum temperature of the warmest month, minimum temperature of the coldest month, mean temperatures of the wettest and driest quarters; precipitation metrics, including the wettest and driest months, seasonality, warmest quarter; and Elevation.

### Biotic variables

The biotic predictors for *M. sexta* and *M. quinquemaculata* included distributions of cultivated host plants, wild host plants, and *C. congregata*. Similarly, the biotic predictors for *C. congregata* included distributions of *M. sexta*, *M. quinquemaculata*, and cultivated and wild host plants. To generate the layer for each biotic predictor, we compiled occurrence records representing known presences of each species. Further, we utilized the disk approach in BIOMOD2 R package, to set bounds and obtain a total of 10,000 pseudo-absences for each species (Barbet-Massin et al. 2012). Pseudo-absences were randomly selected outside the occurrence boundaries, ensuring a buffer of 1 km (∼0.008 degrees) around each presence point (see Kearney et al. 2009) and restricted to regions where the species is unlikely to occur considering optimal present climate conditions. These data points were integrated into an ensemble species distribution model to produce continuous probability maps, where the likelihood of occurrence for each species was represented by values ranging from 0 to 1. The predicted probabilities from the SDMs were subsequently used as inputs (biotic variables) for the target species.

### Species distribution models

We used the R package *sdm* (Naimi and Araújo 2016) to model species distributions (Naimi and Araújo 2016). To overcome the bias and variance across different machine learning algorithms, we used an ensemble approach to combine predictions across three different modelling methods: (a) Generalized Linear Model (GLM), a regression method with linear effects and stepwise selection based on Akaike Information Criteria (AIC); (b) Random Forest model (RF), a machine learning method (default: number of trees = 501; node size = 5); and (c) Maximum Entropy modeling (MaxEnt), a machine learning method with only linear and quadratic features (Marshall et al. 2017; Scriven et al. 2018). GLM algorithms have been successfully used in modeling other insect distributions because they conform well to presence-absence data given their threshold-type response and assume additive or linear relationships for data (Guisan et al., 2002). The RF algorithm is a comparatively robust statistical classifier with high classification accuracy that performs well in modeling complex interactions among predictor variables (Mi et al., 2017). MaxEnt is appealing because it does not require true absences and performs well with minor spatial errors and few occurrence records (Elith et al., 2011).

We modeled species distribution in three categories: abiotic-only model, biotic-only model, and abiotic-biotic model. A randomization approach was used to quantify the relative importance of the abiotic or biotic variable(s) that were most important in predicting where the species were likely to occur under each model category. Correlations between model predictors were analyzed to assess their relationship with species occurrence. Response curves were generated by inserting an “evaluation strip” into the spatial data layers and clipped out to create plots of the modeled responses (Elith et al. 2005). Geographical projections of the models with heat maps were used to visualize the predicted probability of species occurrence in climatic space. All analyses were performed with R version 4.10.0 (R Development Core Team, 2019). Below is a description of the models built for each player in the tri-trophic system.

#### a. Manduca spp

For *M. sexta* and *M. quinquemaculata*, the abiotic-only model incorporated nine abiotic variables that remained after a collinearity check and were considered biologically relevant. The abiotic model and its variables were consistent across species. The biotic-only model incorporated the likelihood of occurrence of *C. congregata*, cultivated and wild host plants. The abiotic-biotic models combined all nine abiotic variables with the likelihood of occurrence of *C. congregata*, cultivated and wild host plants.

#### b. C. congregata

For *C. congregata*, the abiotic-only model included the same nine environmental variables used in the abiotic models for *Manduca* species. The biotic-only model incorporated the likelihood of occurrence for *M. sexta*, *M. quinquemaculata*, and their associated cultivated and wild host plants. The integrated abiotic-biotic model combined all abiotic variables with the predictors from the biotic-only model, to evaluate how abiotic factors, and the distributions of Manduca spp. and its host plants jointly influence the occurrence of *C. congregata*.

#### c. Host plants

Abiotic-only models were developed for both cultivated and wild host plants, each incorporating the same nine abiotic variables. Biotic-only and abiotic-biotic models were not considered for host plants due to their broad ecological niches and to simplify model composition.

### Model evaluation

Model performance was assessed using the Area Under Curve (AUC) and True Skill Statistics (TSS) model evaluation methods based on a confusion matrix (Allouche et al. 2006; Philips et al. 2006). AUC is a measure of rank-correlation; a high AUC means that areas predicted to have a high probability of occurrence tend to be areas of known presence, and locations with lower probability of occurrence prediction values tend to be areas where the species is absent. An AUC score of 0.5 means that the model is as valid as randomly taking a guess, while AUC values >0.8 indicate satisfactory model performance (Philips et al. 2006). TSS relates to the sum of the proportion of presences correctly predicted, and the proportion of absences correctly predicted minus one. Values range from −1 to 1, where a TSS close to 1 means the model has a strong ability to distinguish between areas where the species is present and where it is absent, a TSS of 0 suggests no better performance than random chance, and TSS values less than 0 indicate that the model performs worse than random guessing (Allouche et al. 2006). Both AUC and TSS have been demonstrated to be accurate measures of model performances (Allouche et al. 2006; Philips et al. 2006).

We evaluated the performance of each model category i.e., abiotic-only, biotic-only and/or abiotic-biotic models for Manduca spp., and its parasitoid *C. congregata*. First, we calculated the AUC and TSS values for each statistical algorithm (RF, GLM, and Maxent) under the individual model categories. These categories included abiotic-only, biotic-only, and abiotic-biotic models. The average AUC and TSS for each model category were determined by computing the mean of the AUC values obtained from the three algorithms (RF, GLM, and Maxent). To assess differences in model predictability across the model categories for each species, we performed a one-way ANOVA, comparing the mean AUC and TSS values among the categories.

## Results

### a. Manduca spp

The abiotic model consisted of nine abiotic variables, including maximum temperature of the warmest month, minimum temperature of the coldest month, mean temperatures of the wettest and driest quarters; precipitation of the wettest and driest months, precipitation seasonality, precipitation of warmest quarter; and elevation. Under the abiotic model, minimum temperature of the coldest month was the most important variable for predicting the probability of occurrence for both *M. sexta* and *M. quinquemaculata* (Figure 3a). The probability of *M. sexta* and *M. quinquemaculata* occurrences increased in regions with warmer coldest month temperatures >10 °C (Figure 4a,b). Among the precipitation-related variables, precipitation during the driest month were the strongest predictors of *M. sexta* distribution and precipitation of warmest quarter (summer) was the strongest predictor of *M. quinquemaculata* distribution (Figure 3a). Regions with precipitation > 400 mm in the driest month were suitable for both *M. sexta* and *M. quinquemaculata* occurrences. *Manduca sexta* was found in areas with higher summer precipitation > 400 mm, while *M. quinquemaculata* was less likely to occur in such regions (Figure 4a, b). Further, the probability of both *M. sexta* and *M. quinquemaculata* occurrences increased at elevations below 500m and above 1000 m elevations (Figure 5).

**Figure 3:**
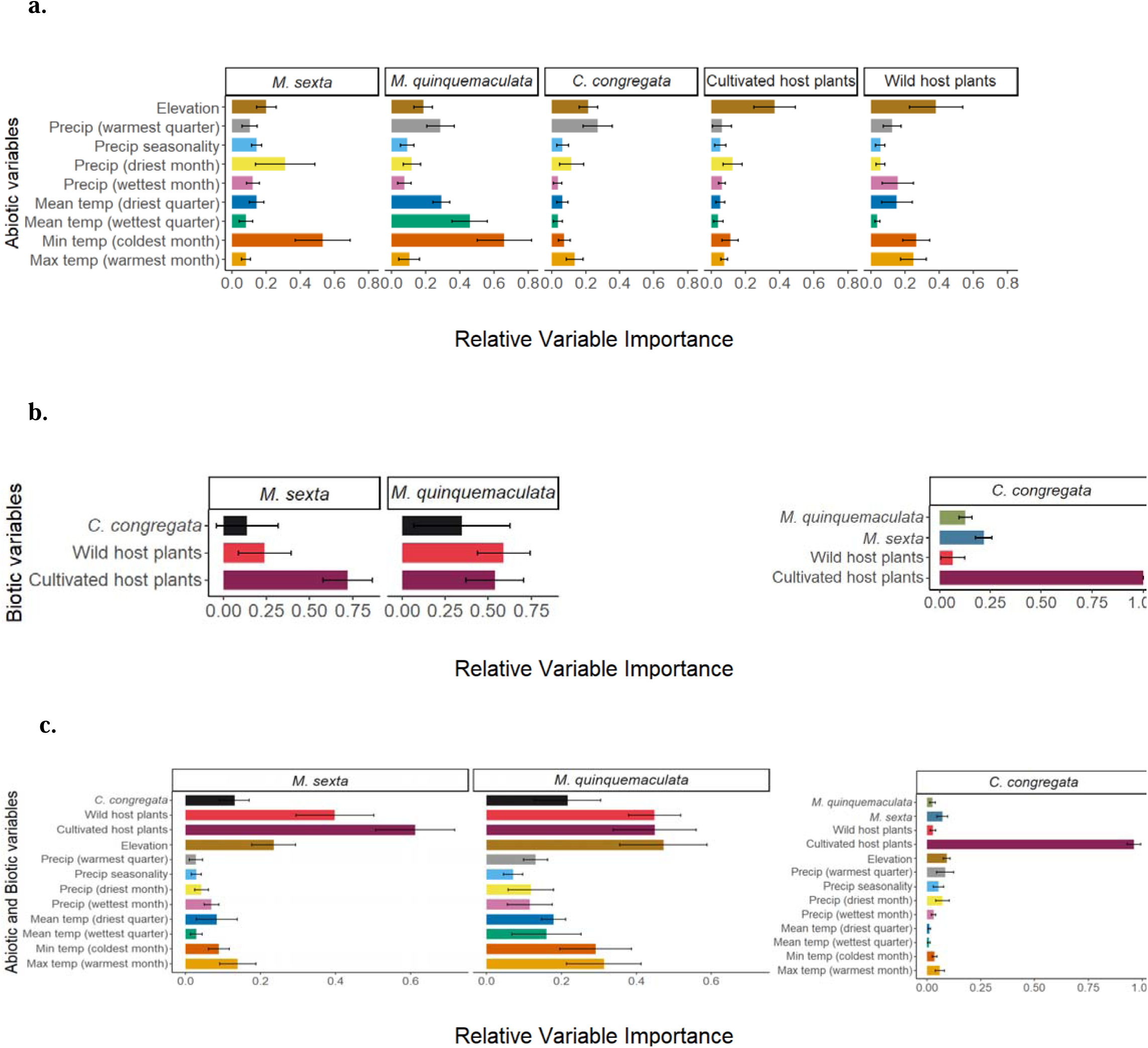
Relative variable importance for the predicted distributions of the *M. sexta* and *M. quinquemaculata*, the parasitoid *C. congregata*, and both wild and cultivated host plants. The y-axis displays the variables included in each model category for each species. (a) results from the abiotic-only model, (b) results from the biotic-only model, and (c) results from the abiotic-biotic model. Bars show means ± 95% CI.

**Figure 4:**
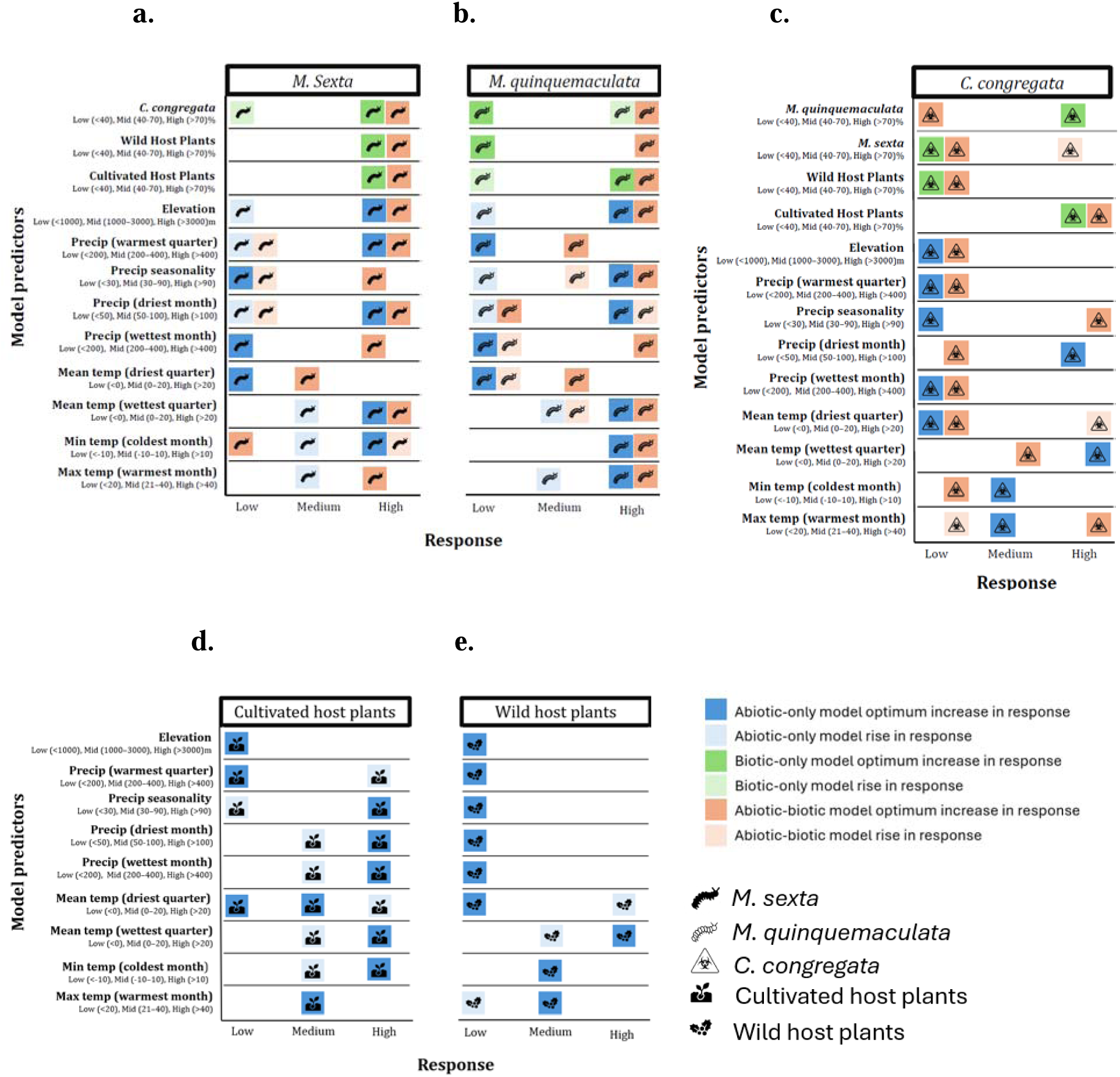
Each panel illustrates how species respond in terms of likelihood of occurrence under low, medium, and high levels of model predictors. Species are represented by unique icons, while model categories—abiotic-only, biotic-only, and abiotic-biotic—are distinguished by unique colors. Lighter shades indicate conditions where the species showed an increase in likelihood of occurrence, while deeper shades represent the conditions associated with the optimum likelihood of occurrence. Panels display results for (a) *Manduca sexta*, (b) *Manduca quinquemaculata*, (c) *Cotesia congregata*, (d) cultivated host plants, and (e) wild host plants.

**Figure 5.**
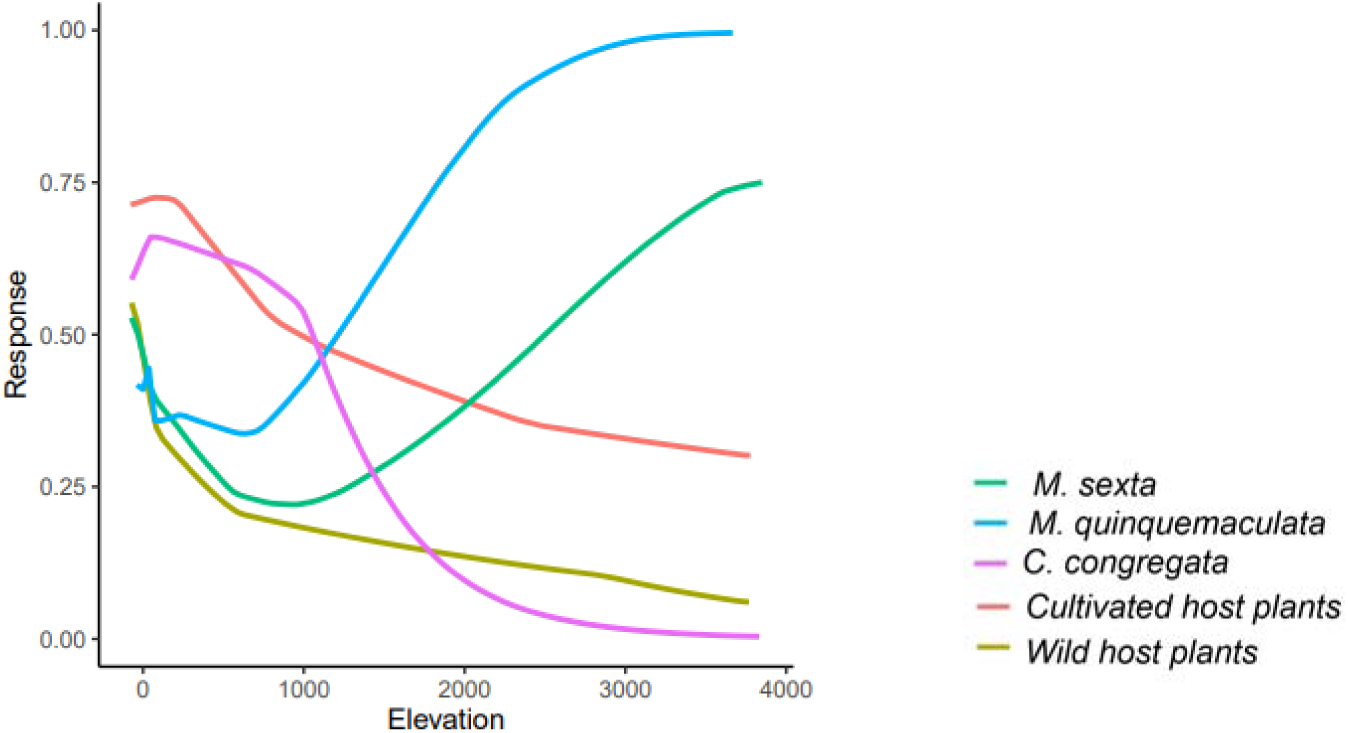
Response curves showing the probability of occurrence for insect hosts (*Manduca sexta* and *Manduca quinquemaculata*), the parasitoid (*Cotesia congregata*), wild host plants (*Solanum dulcamara*, *Nicotiana glauca*, *Datura wrightii*), and cultivated host plants (*Solanum lycopersicum*, *Solanum tuberosum*, *Nicotiana tabacum*) across an elevation gradient. The x-axi represents elevation, while the y-axis represents the probability of occurrence of species in response to elevation.

In the biotic-only model, which examined the impact of cultivated and wild host plants and parasitoid *C. congregata* on the distribution of the two *Manduca spp.*, cultivated host plants emerged as the most influential predictor (Figure 3b). For *M. quinquemaculata*, the likelihood of occurrence of wild host plants and *C. congregata* were equally important predictors. Both herbivores exhibited similar responses to the probability of cultivated host plant occurrences. Specifically, regions with a higher likelihood of *Manduca* spp. presence was correlated with an increased probability of cultivated host plant occurrences (Figure 4a,b). Also, the optimal likelihood of *M. sexta* occurrence was observed in regions with a higher probability of *C. congregata* and wild host plant occurrences, whereas *M. quinquemaculata* did not display a similar pattern (Figure 4a,b).

The most significant predictor in the abiotic-biotic-only model for the two *Manduca spp.* did not change from the biotic-only model results (Figure 3c). However, unlike the biotic-only model, both *M. sexta* and *M. quinquemaculata* showed similar responses to the distributions of host plants and *C. congregata*. The optimal likelihood of occurrence for both *Manduca spp.* Was found in regions with greater than 70% likelihood of occurrence of cultivated and wild host plants, as well as the presence of *C. congregata*. (Figure 4a,b).

### b. C. congregata

Under the abiotic-only model, summer precipitation was the primary abiotic predictor of *C. congregata* distribution, with occurrences positively correlated with increasing levels of summer rainfall, similar to its host *M. sexta* (Figure 3a). Elevation was the next strongest predictor of *C. congregata* distribution. The likelihood of occurrence increased at elevation <500 however unlike its host *Manduca spp.*, at elevations exceeding 1000m, the probability of parasitoid occurrence markedly decreased (Figure 5).

In the biotic-only model, cultivated host plants were the most important biotic predictor of *C. congregata* distribution (Figure 3a). The parasitoid exhibited a greater likelihood of occurrence in regions with a higher probability of cultivated host plant presence. *C. congregata* showed a higher likelihood of occurrence in areas where *M. quinquemaculata* was likely to be present, but not in regions associated with *M. sexta*. wild host plants were generally the least important predictor of *C. congregata* distribution.

The most significant predictor in the abiotic-biotic-only model for the parasitoid did not change from the biotic-only model results (Figure 3c). Cultivated host plants were the most important predictor of occurrence, followed by the host herbivore, *M. sexta*. The parasitoid was likely to occur in regions where cultivated host plants were likely present (Figure 4c). In contrast to the biotic-only models, which indicated that the parasitoid was more likely to occur in areas where *M. quinquemaculata* was present, the abiotic-biotic model suggested that the parasitoid was less likely to occur in areas where *M. quinquemaculata* was found but could co-occur with *M. sexta* (Figure 4c).

### c. Host plants

The results from the abiotic-only model showed the most important abiotic predictor for both the cultivated and wild host plants was elevation (Figure 3a). The probability of occurrence of both types of host plants declined with increasing elevation (Figure 5). The most important temperature-related variable for cultivated and wild host plants was the minimum temperature of the coldest month (Figure 3a). The probability of occurrences of wild and cultivated host plants declined in areas of low minimum temperature <-10°C of the coldest month. Among the precipitation-related variables, precipitation of the driest month was the strongest for cultivated host plants while precipitation of the wettest month was the strongest predictor for wild host plants. Regarding driest month precipitation, both types of host plants occurred where precipitation was <100mm albeit cultivated host plants mostly occurred in regions with relatively wetter summers than wild host plants. Cultivated host plants were also likely to occur in regions with high precipitation of the wettest month but not wild host plants (Figure 4d,e).

### d. Comparison between joint and separated abiotic and biotic models

For *M. sexta*, the mean AUC values for the abiotic-only, biotic-only, and abiotic-biotic models were 0.857, 0.85, and 0.90, respectively. The one-way ANOVA showed no statistically significant differences among the model categories (*F* (2, 6) = 0.689, *p* = 0.538). Similarly, the mean TSS values for these models were 0.577, 0.557, and 0.637, respectively, with no significant differences observed (*F*(2, 6) = 0.341, *p* = 0.724). These findings indicate that the performance of the different model categories was comparable for *M. sexta* in terms of both AUC and TSS.

For *M. quinquemaculata*, the mean AUC values for the abiotic-only, biotic-only, and abiotic-biotic models were 0.817, 0.733, and 0.85, respectively (*F*(2, 6) = 1.403, *p* = 0.316). Corresponding TSS values were 0.513, 0.39, and 0.557 (*F*(2, 6) = 0.955, *p* = 0.436). No statistically significant differences were found among the model categories, consistent with the findings for *M. sexta*.

For *C. congregata*, the mean AUC values were 0.813 for the abiotic-only model, 0.89 for the biotic-only model, and 0.91 for the abiotic-biotic model (*F*(2, 6) = 2.204, *p* = 0.192). The corresponding TSS values were 0.493, 0.63, and 0.677 (*F*(2, 6) = 2.384, *p* = 0.173).

Across all three species, the abiotic-biotic models tended to exhibit higher mean values. However, the differences were not statistically significant.

## Discussion

We employed an ensemble approach to examine the influence of abiotic, biotic, and combined abiotic-biotic factors on the distribution of *M. sexta*, *M. quinquemaculata*, and their shared and sole hymenopterous larval endoparasitoid, *C. congregata*. Additionally, we assessed how abiotic factors alone affected the distribution of cultivated and wild host plants. The difference in predictive ability for abiotic-only, biotic-only, and abiotic-biotic model categories was further analyzed to address how the inclusion of biotic factors alters traditional, abiotic-only SDMs. Our results indicate that in the abiotic-only models, minimum temperature of the coldest month emerged as the most influential predictor for the distribution of both hornworm species, while summer precipitation was the key abiotic factor influencing the distribution of *C. congregata*. However, when biotic variables were included, the presence of cultivated host plants became the primary driver of the distributions of both hornworm species and their parasitoid, *C. congregata*. Our findings revealed no significant difference in the predictive power of models based on isolated or combined abiotic and biotic factors.

Warmer winter temperatures are associated with over-wintering survivorship for herbivore species that overwinter as pupae (Abarca et al., 2024; Crozier, 2003; Mironidis et al., 2010; Morey et al., 2012), such as *M. sexta* and *M. quinquemaculata*. The probability of *M. sexta* and *M. quinquemaculata* occurrences increased in regions with warmer coldest month temperatures, suggesting that their persistence is particularly sensitive to extreme cold. This response to winter temperatures has been shown in other lepidopterans (Crozier, 2003; Pollard, 1988). Previous studies on *Atalopedes campestris* (Sachem Skipper) showed similar constraints where cold tolerance played a pivotal role in its range expansion (Crozier, 2003), and warmer winters facilitated their spread into regions that were previously too cold for overwintering success. Likewise, studies on *Helicoverpa armigera* (cotton bollworm) in Europe and North America demonstrated that cold winter temperatures limited overwintering survival, reducing their capacity to expand into new regions (Mironidis et al., 2010).

Climate-related factors can drive speciation and ecological adaptations in closely related species. An example of sister species that have adapted to different precipitation or climatic conditions is *Odontamblyopus lacepedii* and *Odontamblyopus rebecca*, two eel-goby species. Whereas, *O. lacepedii* is found in the northern, cooler regions, *O. rebecca* occupies southern, warmer areas. This differentiation has been studied through genome sequencing, revealing substantial genetic differences shaped by their respective environmental conditions (Lü et al., 2023). The responses of the two species to summer precipitation diverged: *M. sexta* was found in areas with high summer rainfall (> 400 mm), whereas *M. quinquemaculata* showed a preference for regions with comparatively lower levels of summer precipitation (<200 mm). For *M. sexta*, higher summer rainfall might be beneficial for larval development, potentially due to a denser growth of host plants (Zeppel et al., 2014). In contrast, *M. quinquemaculata* appears to favor drier conditions, which could be linked to its tolerance of different host plants (Fang et al., 2005) or adaptation to less humid microhabitats.

The spatial separation between *M. sexta* and *M. quinquemaculata* in areas with differing precipitation levels may also impact their interactions with *C. congregata*. Occurrences *of C. congregata* correlated positively with increasing levels of summer precipitation, suggesting that areas with higher rainfall provide suitable environmental conditions for the parasitoid and host populations of *M. sexta*. Since *C. congregata* is closely associated with *M. sexta*, its distribution aligns more with regions of higher summer precipitation, reflecting the overlap with its primary host. Thus, variations in precipitation may influence the ecological dynamics between the parasitoid and its hosts, leading to spatially distinct patterns of co-occurrence and niche partitioning.

Elevation emerged as a key abiotic predictor for the distribution of both cultivated and wild host plant groups, with the probability of their occurrence decreasing significantly at higher altitudes. This suggests that the most suitable habitats for these plants are found at lower elevations, where conditions like warmer temperatures and higher soil moisture levels may better support their growth. Additionally, while both host plant types could tolerate dry conditions, cultivated host plants were more commonly found in regions with relatively wetter summers compared to wild host plants. This could be because cultivated host plants are also irrigated or that cultivated species have higher moisture requirements during the growing season, possibly due to their domestication and selection for environments that favor productivity under more consistent moisture availability.

The presence of cultivated host plants emerged as the most important biotic factor, driving the occurrence patterns of both *M. sexta* and *M. quinquemaculata*, as well as *C. congregata*. The strong influence of cultivated host plants highlights the role of agricultural landscapes in shaping the distribution of these species. Since cultivated host plants provide abundant food resources, they likely facilitate higher densities of hornworm populations, which, in turn, sustain larger populations of *C. congregata*. This result aligns with previous studies showing that domesticated plant species can serve as key hotspots for insect populations, including both herbivores and their natural enemies (Garvey et al., 2020). The presence of cultivated host plants may create a resource-rich environment, allowing *M. sexta* and *M. quinquemaculata* to establish and maintain stable populations, while also supporting the reproductive success of *C. congregata*.

Interestingly, although the abiotic-biotic model exhibited consistently higher predictive ability, AUC and TSS model evaluation results did not significantly differ from the abiotic-only or biotic-only models. These findings underscore that while the integrated abiotic-biotic model provides a more comprehensive understanding of species distributions by accounting for both environmental and biological factors, the contribution of these additional biotic predictors may not always be substantial enough to significantly outperform the variation abiotic conditions alone capture. From previous studies, Leach et al. (2016) showed that models with both abiotic and biotic factors (co-occurring species) were better predictors of lagomorph distributions in Europe compared to abiotic-only models. In contrast, Giannini et al. (2013) showed that the inclusion of a biotic interaction for a host-parasitoid system (bumblebees and brood parasites) did not significantly improve the predictive ability of distribution models compared to abiotic-only models; however, in the same study the inclusion of a biotic interaction for a specialist plant-pollinator system (Centris bees and Krameria flowers) did increase model predictive ability. As climate variability continues to impact ecosystems, more comprehensive approaches will be essential for predicting species responses and formulating effective conservation strategies.

One potential explanation for our outcome is the method used to derive the absence data. Absences were generated using pseudo-absence points limited to areas where the species is unlikely to occur, given optimal current climate conditions. This approach, while commonly used (Golicher et al., 2012), may have inadvertently inflated the influence of abiotic factors. Because pseudo-absence data are generated based on climate suitability, abiotic conditions may dominate the model’s predictive capacity, potentially overshadowing the contribution of biotic interactions, even in the combined model. This methodological limitation highlights an inherent challenge in species distribution modeling: the use of pseudo-absences can skew results by disproportionately emphasizing environmental variables. True verified absence data would offer a more accurate baseline for evaluating the roles of both abiotic and biotic factors.

The equivalence in predictive ability of abiotic-only, biotic-only, and abiotic-biotic models may reflect the underlying influence of abiotic factors on the biotic factors. Abiotic factors have indirect effects on biotic interactions, making it difficult to infer the importance of each variable separately for species distributions (Godsoe et al. 2016). Godsoe et al. (2016) simulated the effects of competition and showed that abiotic factors remained strong predictors in SDMs even when competition was known to influence distributions. The prevalence of these indirect effects on SDMs is not well known, but the results here and in other studies (Giannini et al. 2013, Godsoe et al. 2016) emphasize the importance of natural history for SDMs. Research that combines simulation modeling with experimental evidence, greater occurrence records, known absences, and consideration of limitations will be important advances for SDMs. For example, simulation models or process-based models could incorporate data from laboratory and field studies on physiology, development under different climatic conditions, or thermal tolerances for understanding the direct and indirect effects of abiotic and biotic factors on species distributions.

