## Supplementary for "Including biotic factors does not improve species distribution models across three trophic levels"

**Table 1** List of abiotic variables used in the model with their abbreviation and calculated variance inflation factor (VIF).

| Abiotic predictors | VIF |
| --- | --- |
| Maximum Temperature of Warmest Month | 5.850 |
| Minimum Temperature of Coldest Month |  |
| Mean Temperature of Wettest Quarter | 3.282 |
| Mean Temperature of Driest Quarter | 5.416 |
| Precipitation of wettest month | 3.790 |
| Precipitation of driest month | 9.064 |
| Precipitation seasonality | 5.104 |
| Precipitation of warmest quarter | 5.700 |
| Elevation | 5.695 |

These climate variables, elevation and the biotic predictors were used in the SDM to characterize the location of suitable biological-climate space for the hornworms under historical climate scenarios.

| a. *M. sexta* | b. *M. quinquemaculata* | c. *C. congregata* |
| --- | --- | --- |
| 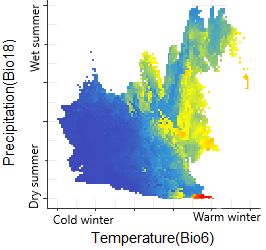 | 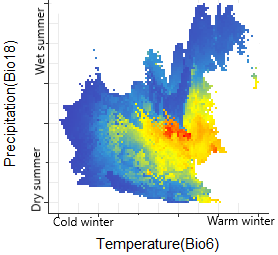 | 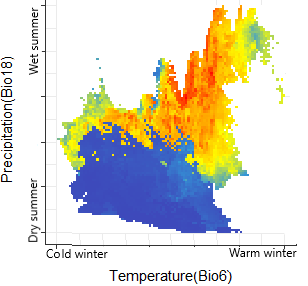 |
| d. Cultivated Host plants | e. Wild Host plants |  |
| 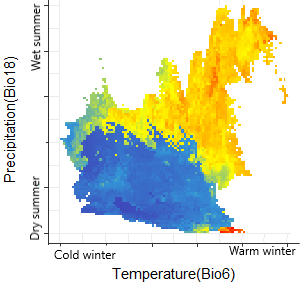 | 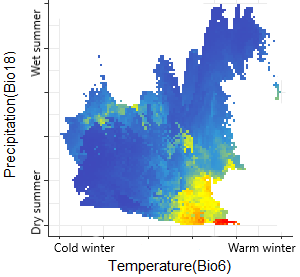 | 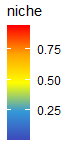 |

Figure 3. Two-dimensional visualization of the ecological niche of M. sexta, M. quinquemaculata, parasitoid *C. congregata*, and cultivated (CHP) and wild (WHP) host plants for precipitation of the warmest quarter (Bio18) and minimum temperature of the coldest month (Bio 6). The combinations predicted habitable for each species are shown as yellow-red. Mean temperature and precipitation ranged between -24.4 to 18.5°C and 2 to 647mm, respectively.
